# Protein Lactylation and Metabolic Regulation of the Zoonotic Parasite *Toxoplasma gondii*

**DOI:** 10.1101/2022.05.19.492655

**Authors:** Deqi Yin, Ning Jiang, Chang Cheng, Xiaoyu Sang, Ying Feng, Ran Chen, Qijun Chen

## Abstract

The biology of *Toxoplasma gondii*, the causative pathogen of one of the most wide-spread parasitic diseases remains poorly understood. Lactate, which is derived from glucose metabolic pathways, is considered to be not only an energy source in a variety of organisms including *Toxoplasma gondii*, but also a regulatory molecule that participates in gene activation and protein functioning. Lysine lactylation is a type of posttranslational modifications (PTMs) that was recently associated with chromatin remodeling, but lysine lactylation of histone and non-histone proteins has not yet been studied in *T. gondii*. To examine the prevalence and function of lactylation in *T. gondii* parasites, we mapped the lactylome of proliferating tachyzoite cells and found 1964 lactylation sites on 955 proteins in the *T. gondii* RH strain. The lactylated proteins were distributed in multiple subcellular compartments and were closely related to a wide variety of biological processes, including mRNA splicing, glycolysis, aminoacyl-tRNA biosynthesis, RNA transport, and multiple signaling pathways. We also performed chromatin immunoprecipitation sequencing (ChIP-seq) analysis with a lactylation specific antibody, the results revealed that histone H4K12la and H3K14la were enriched in the promoter and exon regions of *Toxoplasma gondii* genes associated with microtubule-based movement and cell invasion. We further confirmed the de-lactylase activity of histone deacetylase TgHDACs 2, 3, and 4, and that treatment with anti-histone acetyltransferase (TgMYST-A) antibodies profoundly reduced protein lactylation in the parasites. This study offers the first dataset of the global lactylation proteome and provides a basis for further dissection of the functional biology of *Toxoplasma gondii*.

## Introduction

*Toxoplasma gondii* is an exclusively intracellular apicomplexan parasite that infects all nucleated vertebrate cells and a globally prevalent protozoan parasite that potentially infects more than 30% of the world’s population [1]. This parasite has become a major opportunistic pathogen that causes toxoplasmosis, which is considered one of the most widespread zoonotic diseases [2]. *T. gondii* has a complicated life cycle, which mainly involves the sexual and asexual phases, and primarily exists in two infectious forms, including rapidly replicating tachyzoites and slowly dividing bradyzoites in the tissues of its intermediate hosts (warm-blooded animals) [3]. Although the parasite typically causes mild symptoms in immunocompetent individuals, the developing fetuses, and immunocompromised individuals (particularly acquired immune deficiency syndrome patients) do develop severe symptoms that might even be life threatening [4].

After invading a host cell, *T. gondii* disseminates in the form of rapidly replicating haploid tachyzoites throughout the nucleated cells in the host. In certain organs and tissue types, tachyzoites transform into bradyzoites to form cysts (a semidormant form associated with slow replication) and thereby establish a continuously latent infection state in the host. Once the immune capacity of the host is weakened, the bradyzoites are activated and transformed into tachyzoites to re-infect cells [5]. The sexual reproduction of *T. gondii* takes place in the intestinal epithelial cells of felines (the terminal hosts), and the parasites in the form of oocysts are released with feces, and thus contaminate the environment. Humans and warm-blooded animals are typically infected by exposure to tissue cyst- or oocyst-contaminated food or water. At present, the effective anti-*T. gondii* infection agents are restricted to some common metabolic inhibitors, including azithromycin, pyrimethamine, spiramycin, sulfonamides, trimethoprim-butoxypyrimidine, and clindamycin [6]. Notably, these agents share a common limitation in that they are only effective on tachyzoites at the replication phase and exert only a slight or even no effect on semidormant bradyzoites, and some might even cause severe cytotoxicity to the host. In this regard, further knowledge regarding the biological characteristics of *T. gondii* is warranted for the discovery of new drug targets.

Type I *T. gondii* strain (such as the RH strain) is highly virulent and lethal to mice, resulting in severe clinical manifestations. However, type II strain (such as the ME49 strain) exhibit relatively low pathogenicity to murine hosts, typically manifesting as chronic infections [7]. The proliferation of *T. gondii* is strictly dependent on precise gene expression control that ensures appropriate protein profiles [4]. Posttranslational modifications (PTMs) constitute one of the critical response mechanisms of *T. gondii* to external environmental stimulation and may control its lifecycle transitions [8]. PTMs, which involve the process of adding chemical moieties to one or more amino acid residues for the regulation of protein function, affect almost all aspects of pathogenesis and cell biology of the parasite [5]. Specifically, these modifications affect the spatial conformation, subcellular localization, folding and stability of proteins, and protein–protein interactions (PPIs) by changing the physicochemical properties of proteins, and these effects significantly regulate the complex life activities of the parasites. Therefore, revealing the regulatory mechanism of PTMs in *T. gondii* will promote the discovery of parasite-specific means for toxoplasmosis treatment and prevention. To date, more than 12 types of PTMs, namely, 2-hydroxyisobutyrylation, malonylation, acetylation, succinylation, crotonylation, ubiquitination, methylation, palmitoylation, N-myristoylation, SUMOylation, phosphorylation, and O-GlcNAcylation, have been discovered in *T. gondii* [9–15]. Among these PTMs, phosphorylation, which is a highly dynamic type that plays a vital role in cell signal transduction, represents the most common PTM in *T. gondii* [16]. We have previously revealed that 2-hydroxyisobutyrylation covers approximately 23.2% and 20.5% of the proteomes of *T. gondii* RH and ME49 strains, respectively, and plays an essential role in gene expression, energy metabolism, and invasion [15]. In addition, 1061 crotonylated proteins in the *T. gondii* RH strain and 984 in the ME49 strain have also been detected, which are strictly related to energy metabolism, invasion, and protein synthesis and degradation [15]. Lysine acetylation, which covers approximately 5% of the predicted proteome of *T. gondii*, is associated with protein translation and metabolism [17, 18]. Ubiquitination covers up to 5% of the *T. gondii* proteome and is considered a key regulator of cell cycle transition and cell division [19]. Furthermore, 370 arginine monomethylated proteins, which cover approximately 4.5% of the proteome, are involved in transcriptional regulation, DNA repair, and RNA metabolism [10]. Collectively, PTMs display high abundance in *T. gondii* and play an integral role in multiple vital biological pathways.

Lactate was originally thought to be a metabolite of glycolysis and an energy source [20, 21]. Lysine lactylation (Kla), as a novel histone marker associated with gene activation, can be obtained from the cellular metabolite lactate and detected through a + 86.03 Da mass shift of lysine residues [22]. Lactate is currently considered a precursor that stimulates Kla and is produced via the glycolytic pathway.

In this study, we leveraged the first comprehensive analysis of lactylome of *T. gondii* and dissected the regulatory potential of the Kla, which opened a new venue for elucidation of the parasite biology and digging novel parasiticide targets.

## Results

### Identification of Kla in the proteome of the *T. gondii* RH strain

To characterize the global Kla profile of *T. gondii*, whole tachyzoite lysates of the parasites were analyzed by LC-MS/MS (Figure S1A). The protein lactylation in tachyzoites was then verified by Western blotting using a pan anti-Kla antibody. Multiple proteins were recognized (Figure 1A) by the antibody. Presence of lactylated proteins in tachyzoites of *T. gondii* RH and ME49 strains were also confirmed with an immunofluorescence assay (IFA) with anti-L-lactyllysine antibody (Figure 1B and C). The results showed that proteins with lactyllysine were widely distributed inside multiple discrete compartments in *T. gondii*.

**Figure 1.**
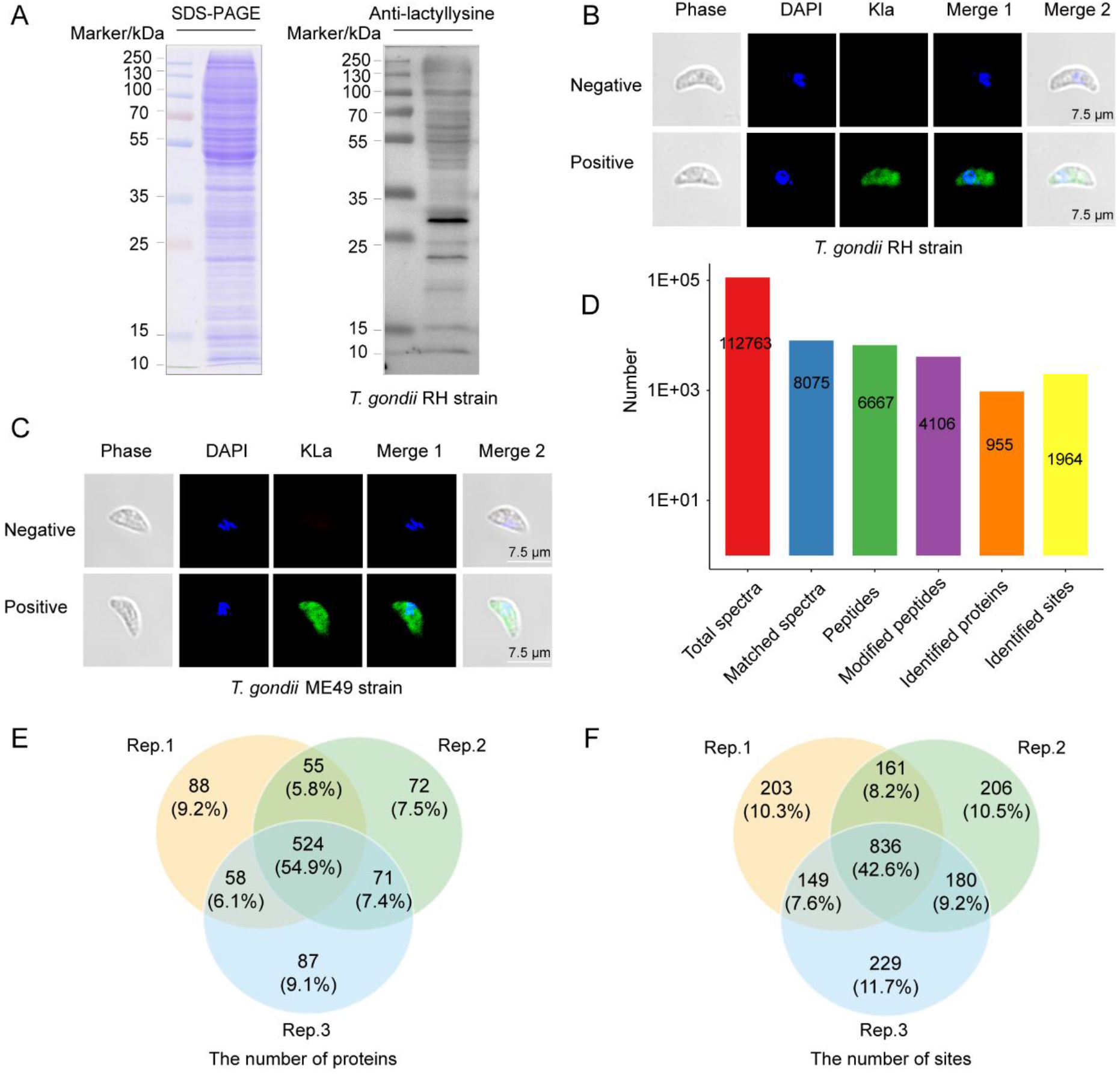
Lactylation is widespread in *Toxoplasma gondii* illustrated by Western blotting, IFA and basic information of LC-MS/MS. **A.** SDS-PAGE and Western blotting analysis of tachyzoite lysate (20 μg) probed with anti-lactyllysine antibody. **B.** IFA of paraformaldehyde-fixed tachyzoites of *T. gondii* RH strain with an anti-lactyllysine antibody (green). The negative control was the paraformaldehyde-fixed tachyzoites of *T. gondii* incubated with rabbit IgG. Nuclei were stained with DAPI (blue). **C.** IFA of tachyzoites of *T. gondii* ME49 strain with an anti-lactyllysine antibody (green). The negative control was tachyzoites incubated with rabbit immunoglobulin G (IgG). Nuclei were stained with DAPI (blue). **D.** Basic information of LC-MS/MS data of lysine lactylation. **E.** Venn diagram showing the overlapping proteins obtained from triplicate replications. **F.** Venn diagram showing the overlapping lactylation sites obtained from triplicate replications. IFA, immunofluorescence assay; LC-MS/MS, liquid chromatography-tandem mass spectrometry; IgG, immunoglobulin G; SDS-PAGE, sodium dodecyl sulfate polyacrylamide gel electrophoresis.

In total, we identified 1964 Kla sites on 955 proteins in triplicate experiments (Figure 1D–F, Table S1). Additionally, we evaluated the mass error to verify the accuracy of the MS data. The MaxQuant data were filtered according to the following thresholds: mass error ≤ 5 ppm, false discovery rate (FDR) of ≤ 1%, and localization probability > 75%. The results suggested that the mass validation of the large-scale data conformed to the experimental standards (Figure S1B). In addition, the distribution of peptide lengths identified met the quality control requirements (Figure S1C). The majority of lactylated proteins (86.1%) contained one to three Kla sites, whereas approximately 5.4% of the proteins contained more than five Kla sites (Figure S1D). We identified three proteins containing the highest number of Kla sites, including the translation elongation factor 2 family protein (16 sites), heat shock 70 protein (14 sites), and elongation factor 1-alpha (14 sites).

### Lactylated proteins are widely distributed in multiple subcellular compartments in *T.gondii*

To further explore the possible biological functions of the lactylated proteins, the modified proteins were statistically classified into Gene Ontology (GO) secondary annotations, as shown in Figure S2 (Table S2). The lactylated proteins were mainly distributed in intracellular organelle terms, and were associated with various molecular functions, such as protein binding, organic cyclic, and heterocyclic compound binding. The top three biological processes were cellular metabolic process, organic substance metabolic process, and primary metabolic process.

Based on the annotation results, we performed a subcellular localization analysis of the modified proteins, as exhibited in Figure 2A (Table S3). The top three localizations of the lactylated proteins were the nucleus (n = 426, 44.61%), cytoplasm (n = 186, 19.48%), mitochondria (n = 168, 17.59%). The proteins distributed in the plasma membrane and extracellular space were 69 (7.23%) and 55 (5.76%), respectively.

**Figure 2.**
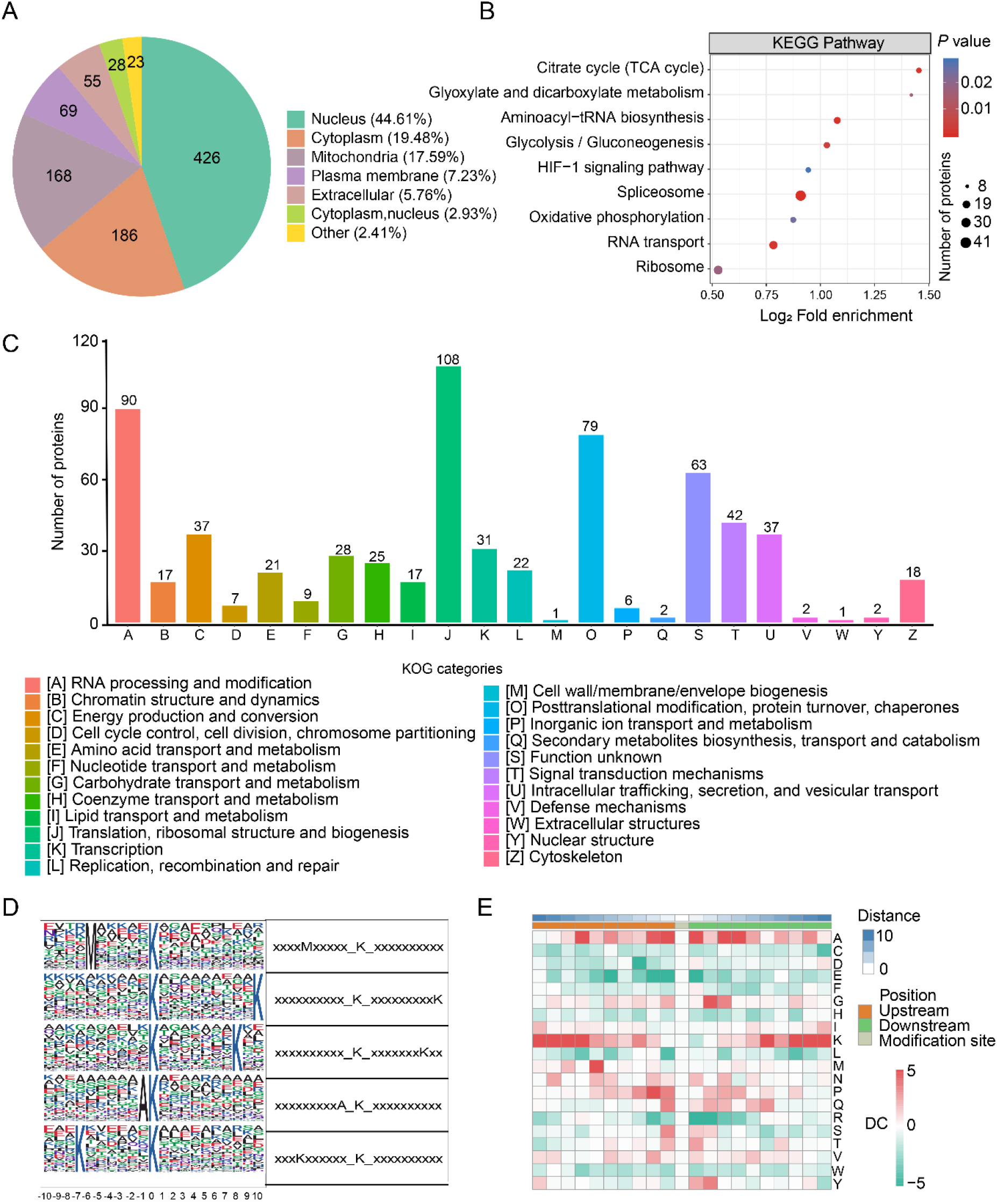
Lactylated proteins associated with a variety of biological processes. **A.** Subcellular localization of the identified lactylated proteins. **B.** Functional KEGG pathway enrichment analysis of lactylated proteins (Fisher’s exact test, *P* < 0.05). **C.** The functional classification of lactylated proteins was performed using the KOG database. The horizontal axis indicates the KOG category, and the ordinate is the number of proteins. **D.** Probability sequence motifs of lactylation sites consisting of 20 residues surrounding the targeted lysine residue were produced using Motif-x. The size of each letter correlates to the frequency of that amino acid residue occurring in that position. **E.** A heat map shows the high (red) or low (green) frequency of occurrence of amino acids at specific positions surrounding of the lactylated lysines. Red box indicates that the amino acid occurring more frequently near the modification site, and green box indicates that the amino acid is less frequent near the modification site. KEGG, Kyoto Encyclopedia of Genes and Genomes; KOG, Eukaryotic Orthologous Groups; DC, difference score.

### Lactylated proteins are enriched with multiple biological function

The biological process of GO category demonstrated that the lactylated proteins were strongly enriched in ribonucleotide monophosphate and triphosphate biosynthetic processes (Figure S3 and Table S4). Furthermore, we also constructed a GO-enriched directed acyclic graph (DAG) of each GO category to provide a visual feature of the lactylated proteins (Figure S4). Notably, mitochondrial components were extremely enriched in the DAG.

Protein domain enrichment revealed that RNA recognition motif was the most highly enriched domain, implying the positive role of lactylation in RNA-binding proteins (RBPs). Furthermore, pyridine nucleotide-disulfide oxidoreductase, elongation factor Tu domain, and the ATP synthase alpha/beta family were also predicted to be significantly enriched (Figure S5 and Table S5).

As revealed by KEGG pathway enrichment, lactylated proteins were mainly enriched in multiple central pathways, including the spliceosome, aminoacyl-tRNA biosynthesis, ribosome, HIF-1 signaling, RNA transport, glycolysis/gluconeogenesis, oxidative phosphorylation, citrate cycle (TCA cycle), and glyoxylate and dicarboxylate metabolism (Figure 2B and Table S6).

Overall Eukaryotic Orthologous Group (KOG) has been developed as a tool for the large-scale functional and evolutionary analysis of proteins. Here, we performed a functional classification analysis of the lactylated proteins based on the KOG database. The top five KOG terms obtained from the analysis of the lactylated proteins were energy production and conversion [C], translation, ribosomal structure and biogenesis [J], signal transduction mechanisms [T], RNA processing and modification [A], and posttranslational modification, protein turnover, and chaperones [O] (Figure 2C and Table S7).

### Lactylation in proteins frequently occurred in peptide regions with conserved motifs

The frequencies of lactylated amino acids (from position −10 to +10) in all identified peptides were statistically analyzed using MoMo software to assess the distinctive characteristics of the sequences surrounding the lactylation sites. According to the enrichment statistics, we retrieved five conserved motifs (Figure 2D). Notably, motifs M-Kla, A-Kla, and K-Kla were quite conserved. We established an enrichment heat map to further analyze the motif sequences (Figure 2E and Table S8). The results showed that methionine (M), alanine (A), and lysine (K) residues often appeared upstream of the lactylation sites and that alanine (A), glycine (G), and lysine (K) residues were more frequently found downstream of the lactylation sites.

### Lactylated proteins are involved in gene transcription and expression

The spliceosome machinery, which is widespread in eukaryotes, is mainly composed of five ribonucleoproteins, namely, U1, U2, U4, U5,and U6 small nuclear RNA (snRNA), and mainly functions to remove transcript regions generated from introns [23]. In this study, 49 lactylated proteins were closely associated with spliceosomes (Figure S6 and Table S9). Notably, 14.29% (7/49) of the proteins contained at least three lactylation sites, and these proteins included DEAD/DEAH box helicase domain-containing protein, small nuclear ribonucleoprotein G, SKIP/SNW domain-containing protein, crooked neck family 1 protein isoform 2, RNA recognition motif-containing protein, pre-mRNA processing splicing factor PRP8 and heat shock protein HSP70. There is evidence that HSP70, may also be involved in the splicing process as a chaperone [24]. Strikingly, the heat shock protein HSP70 was the protein with the highest number of Kla sites (n = 14), which suggested that it might play an essential role in the spliceosome in *T. gondii*.

RBPs play indispensable roles in transcription, translation, and RNA processing in most eukaryotic cells [25–27]. Gene regulation and expression are mainly modulated by transcription-related proteins, such as transcription factors (TFs), splicing factors, RNA polymerases, RBPs, and DNA-binding proteins (DBPs), which are particularly important for the growth and reproduction of *T. gondii*. We summarized the lactylated proteins among the sets of TF-related proteins, RNA polymerase-related proteins, DBPs and RBPs, and found that they are associated with a range of processes including gene transcription, gene splicing, protein translation, and processing (Figure 3 and Table S10).

**Figure 3.**
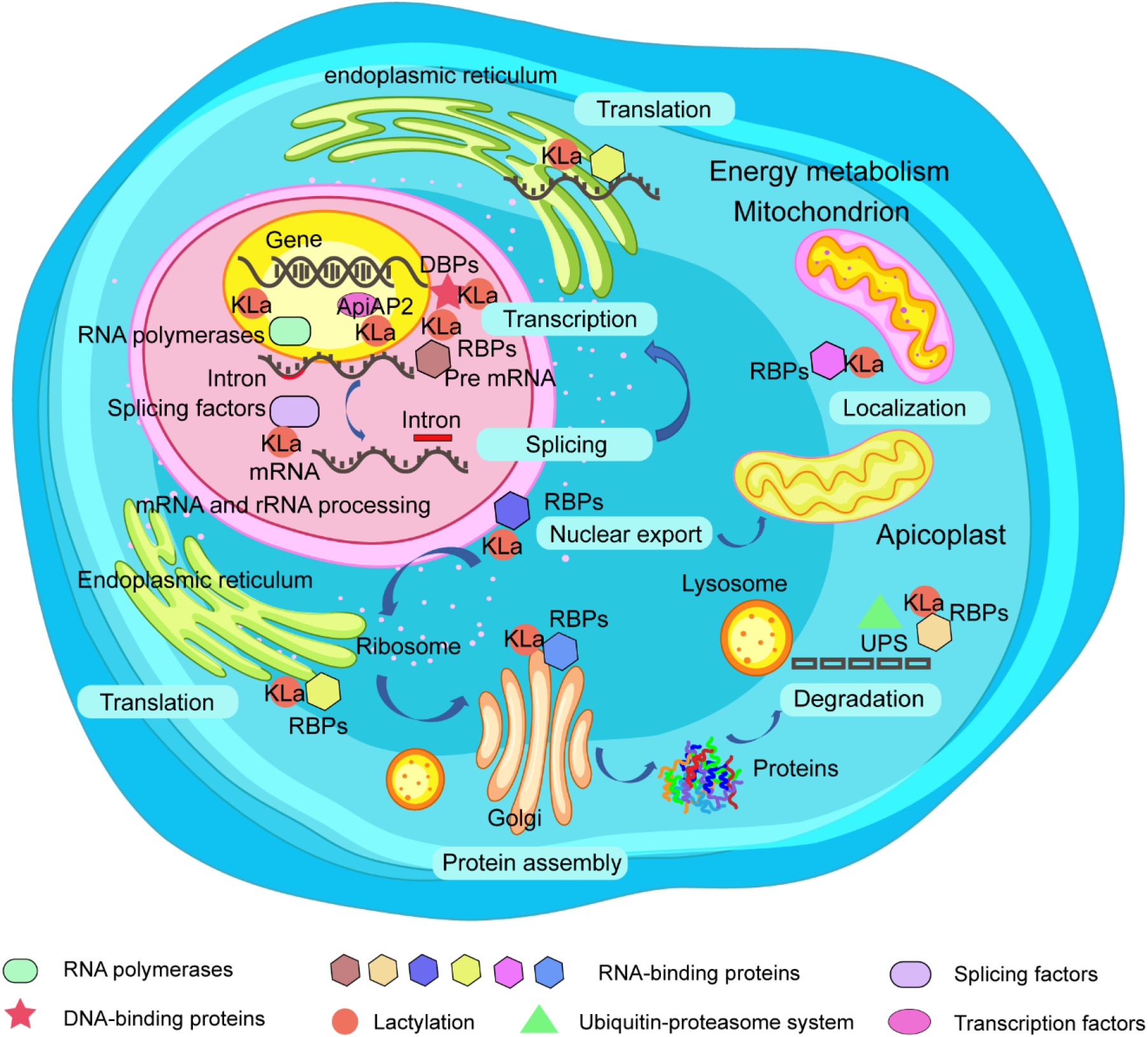
Schematic diagram of the involvement of lactylation in gene regulation. Schematic diagram showing that lactylated proteins such as transcription factor proteins, RNA polymerase-related proteins, DBPs, and RBPs are involved in the transcriptional and translational regulatory processes. The font inside the blue box indicates that lactylated RBPs might play a wide range of roles in various posttranscriptional processes at different cellular locations. Lactylated RBPs and splice factors are major players in the splicing of premRNA into mature mRNA in the nucleus. In addition, lactylated RBPs can assist the translation of mRNA into proteins in the ribosome, and some RBPs might be degraded by the ubiquitin proteasome system. DBPs, DNA binding proteins; RBPs, RNA binding proteins; UPS, ubiquitin-proteasome system.

### Enzymes in glycolysis/gluconeogenesis, the TCA cycle and oxidative phosphorylation were predominantly lactylated

Glycolysis and the TCA cycle have been recognized as the main central carbon metabolic pathways of fast-growing tachyzoites and slowly developing bradyzoites of *T. gondii* [28, 29]. Here, the Kla of key enzymes that mediate the glycolysis/gluconeogenesis and TCA cycle metabolic pathways were thoroughly analyzed (Figure S7). 14 regulatory enzymes related to glycolysis/gluconeogenesis pathway with 43 Kla sites were identified (Table S11), including phosphoglycerate kinase (PGKI), fructose-1,6-bisphosphate aldolase (ALD2), glyceraldehyde-3-phosphate dehydrogenase (GAPDH1), fructose-1,6-bisphosphate aldolase (ALD1), glucosephosphate-mutase (GPM1), fructose-bisphosphate I (FBP 1), fructose-bisphosphate II (FBP 2), enolase 2 (ENO 2), enolase 1 (ENO 1), phosphoglycerate mutase PGMII, pyruvate kinase PyKI, pyruvate kinase PyKII, lactate dehydrogenase (LDH1), and hexokinase (HK). Notably, phosphoglycerate kinase (PGKI) and fructose-1,6-bisphosphate aldolase contained the most Kla sites (n = 6).

For the TCA cycle, a total of 12 enzymes with 47 Kla sites were identified (Table S11), including citrate synthase I (CSI), aconitate hydratase ACN/IRP, isocitrate dehydrogenase (IDH), 2-oxoglutarate dehydrogenase e1 component, putative (2-ODE1), dihydrolipoyllysine-residue succinyltransferase component of oxoglutarate dehydrogenase (DLST), malate dehydrogenase (MDH), succinyl-CoA-synthetase alpha (SCSA), succinate-Coenzyme A ligase, beta subunit, putative (SCAL), succinate dehydrogenase [ubiquinone] iron-sulfur protein (SDIS), flavoprotein subunit of succinate dehydrogenase (FSSD), pyruvate carboxylase (PC), and fumarate hydratase (FH). Notably, aconitate hydratase ACN/IRP was the critical regulatory enzyme in the TCA cycle with the most Kla sites (n = 8). Five Kla sites (K176, K276, K420, K455, and K514) on the pyruvate dehydrogenase complex, which are located in the apicoplast to drive *de novo* fatty acid biosynthesis of the parasite [30], were detected.

In addition, 14 proteins with 27 Kla sites were also associated with oxidative phosphorylation (PATHWAY: map00190) (Table S12), including ubiquinol cytochrome c oxidoreductase, ATP synthase beta subunit ATP-B, type I inorganic pyrophosphatase, ATP synthase F1, delta subunit protein, vacuolar ATP synthase subunit, vacuolar ATP synthase subunit, ATPase synthase subunit alpha (putative), ATP synthase F1 gamma subunit, and copper chaperone COX17-1 (putative).

### Invasion-associated proteins of *T. gondii* were lactylated

Like most protozoan parasites, *T. gondii* utilizes a unique movement machinery involving the actin-myosin complex for gliding to invade host cells [31]. We systemically analyzed the lactylated proteins involved in parasite gliding motility, activation of host cell entry, and egress from infected cells (Table S13). A total of six myosins (myosin A, myosin C, myosin D, myosin F, myosin light chain 2, and myosin head domain-containing protein) containing 19 Kla sites and two actin-related proteins (including actin ACT1 and actin-depolymerizing factor ADF) with seven Kla sites were detected. Furthermore, *T. gondii* typically contains three secreted organelles, namely, micronemes (MICs), rhoptries (RONs and ROPs), and dense granules (GRAs), which are indispensable for the invasion and survival in host cells [32, 33]. We detected various degrees of Kla on MIC1 (K157), MIC2 (K630), RON6 (K11), ROP9 (K79, K138, K144, and K347), ROP18 (K202), GRA12 (K74), and the rhoptry proteins (K954 and K956). Additionally, nine Kla sites were detected on three calcium-dependent protein kinase family proteins (CDPKs), including CDPK1 (K50, K59, K80, K93, and K341), CDPK2A (K165, K729, and K777), and CDPK9 (K5), all these enzymes facilitate parasite gliding motility, cell invasion, egress, intracellular development, and reproduction [34].

### Annotation of lactylated proteins in protein interaction network

A total of 300 modified proteins with 2675 nodes was mapped to the protein interaction database using STRING (confidence score > 0.7), and Cytoscape software was used to obtain a comprehensive view of the interaction network of the lactylated proteins, which were significantly related to various biological pathways, such as ribosome, carbon metabolism, spliceosome, aminoacyl-tRNA biosynthesis, and ribosome biogenesis (Figure S8 and Table S14). The significant hub proteins included bifunctional dihydrofolate reductase-thymidylate synthase (*TGME49_249180*), heat shock protein HSP70 (*TGME49_273760*), Myb family DNA-binding domain-containing protein (*TGME49_275480*), elongation factor 1-gamma, putative (*TGME49_300140*), proliferation-associated protein 2G4 (*TGME49_279390*), and fibrillarin (*TGME49_311430*), which formed a dense protein interaction network.

### Histone lactylation were profoundly identified in *T. gondii*

Histone modification is one of the most important mechanisms in the epigenetic regulation of eukaryotic genes [35]. Here, a comprehensive map of histone PTMs in *T. gondii* was established (Figure 4 and Figure S9). 17 Kla sites on five histones were detected in the *T. gondii* RH strain (Table S15), including H2A.Z (K5, K9, K17, K23, K142, and K150), H2B.Z (K3, K8, K18, and K104), H2Bb (K46 and K98), H3 (K14, K23, K27, and K122), and H4 (K12). And 168 (87.0%) of the PTMs sites were concentrated in H2B.Z, H2A.Z, H2Ba, H3, and H4, and most of the PTM sites were located at the N-termini of histones H2A.Z, H4, H2B.Z, and H3 but not H2Ba (C-terminus). Notably, more than four PTMs were detected in H2BaK70, H2BaK99, H2BaK111, H2B.ZK18, H2A.ZK9, H2A.ZK17, H3K23, H3K27, H3K56, H3K79, H3K122, H4K12, H4K31, and H4K67. Histone modifications are essential in the regulation of processes of gene transcription and expression [36, 37]. However, information regarding H4K12la and H3K14la is still limited. To further confirm the presence of H4K12la and H3K14la, we performed Western blotting and IFA analysis using anti-lactyl-histone H4 (Lys12) and anti-lactyl-histone H3 (Lys14) antibodies, and Western blotting observed that the H4K12la and H3K14la signals co-migrated with bands of approximately 11 kDa and 15 kDa, respectively, which were consistent with the sizes of histones H4 and H3 (Figure 5A). We also confirmed that H4K12la and H3K14la were mainly distributed in the nuclei of the intracellular and extracellular stages by IFA (Figure 5B). The MS/MS spectra of the Kla sites of H3K14 and H4K12 are shown in Figure 5C and D (Table S15).

**Figure 4.**
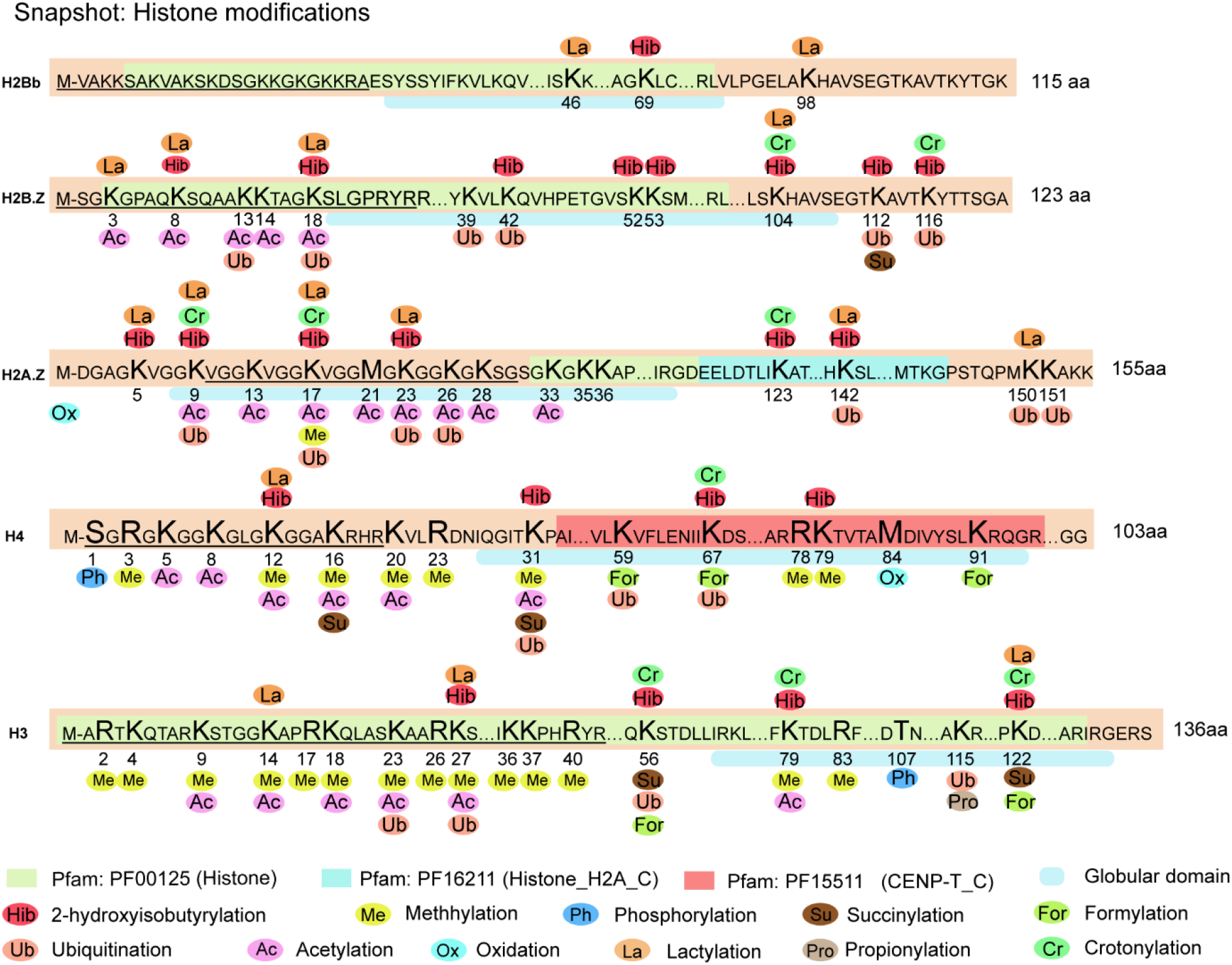
Multiple PTMs occur at the same histone sites. The numbers on the image indicate the specific locations of the modification sites on the histones. Ellipses with different colors represent different types of PTMs. Different colored boxes represent different domains. A horizontal line describes the characteristics of a region (amino acid) in mediating protein-protein interactions or biological processes (Specific information from Universal Protein database, https://www.uniprot.org/uniport). PTM, protein translational modification.

**Figure 5.**
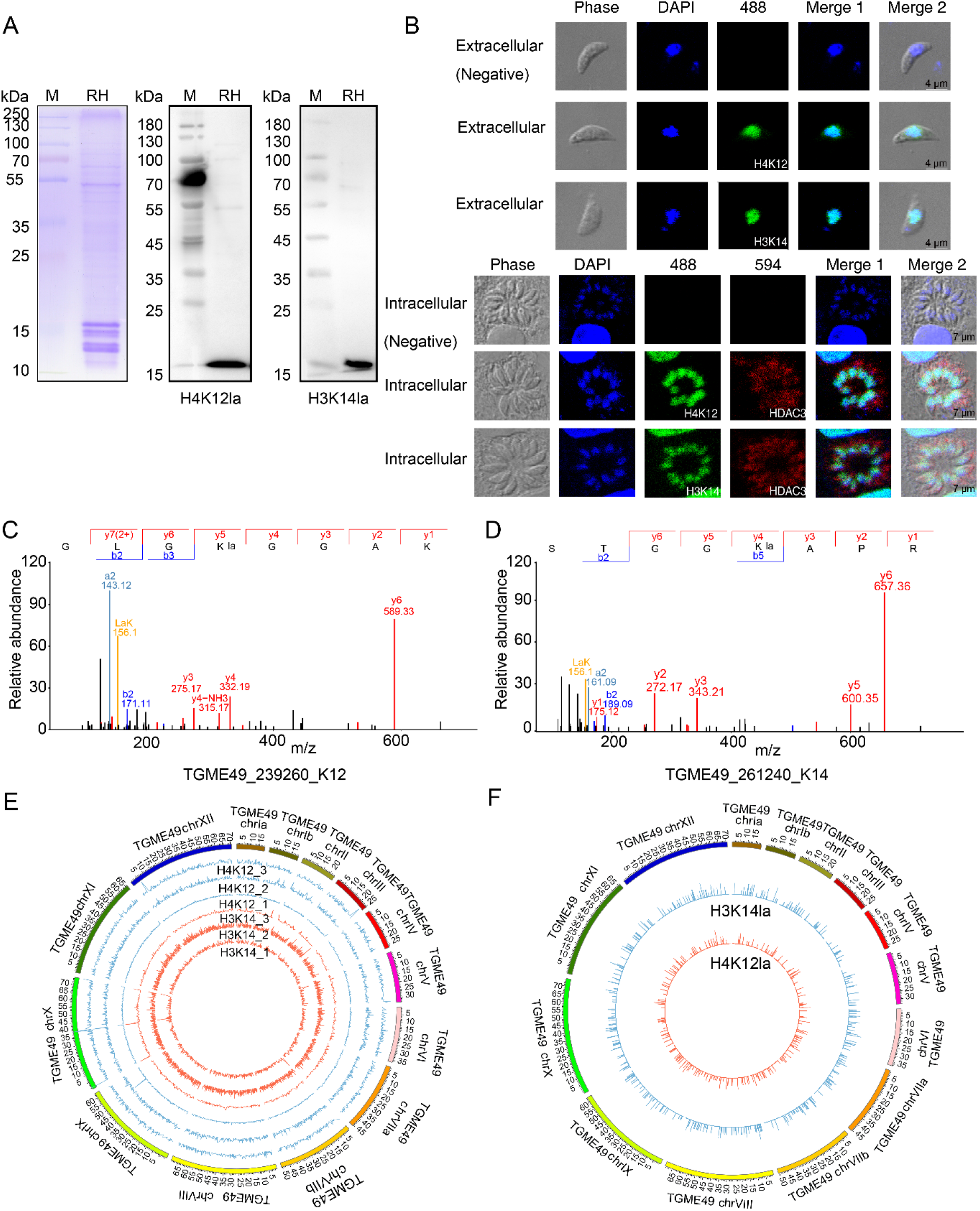
H4K12 and H3K14 of *T. gondii* were identified as histone lactylation sites. **A.** Western blotting of histone H4K12la and H3K14la. **B.** Indirect immunofluorescence assay of histone H4K12la and H3K14la in the intracellular and extracellular stages. Negative control is rabbit IgG. **C.D.** MS/MS spectra of histone H4K12la and H3K14la derived from *T. gondii*,its synthetic counterpart, and their mixture. The a, b ion refers to the N-terminal parts of the peptide, and the y ion refers to the C-terminal parts of the peptide. **E.** Distribution of binding regions of histone lactylation sites (H4K12la and H3K14la) on chromosomes-circos map. The outermost circle shows the chromosomes and the inner circle shows the distribution trend for each sample. Data represent three independent experiments. **F.** Distribution of overlapping peaks of H4K12la and H3K14la in whole genome. The length of the circled bars represents the degree of enrichment, and the position within the circle corresponds to the position of the peaks. ChIP-seq, chromatin immunoprecipitation sequencing.

We next investigated the binding regions of H3K14la and H4K12la on chromosomes by ChIP-seq (Table S16). A total of 1799, 4011, and 1153 peaks were identified in the genomic binding regions of H3K14la in three ChIP-seq experiments. And 2219, 2292, and 2134 peaks were identified in the H4K12la ChIP-seq assay. Three biological replicates of ChIP-seq for H4K12la and H3K14la were performed, respectively, and the distribution of all peaks mapping the genome was shown using a circos plot (Figure 5E and Table S17). Analysis of the average enrichment signal of H4K12la and H3K14la in all gene regions showed that they were mainly distributed around the TSS (Transcriptional Start Site) region (Figure S10A and B). The unique peaks in the binding regions of H4K12la and H3K14la were distributed on all 14 chromosomes of *T. gondii*, with TGME49_chrX, TGME49_chrVIII, and TGME49_chrXII being the most widely distributed (Figure 5F, Figure S10C and D).

The genomic distribution of H3K14la and H4K12la enriched regions can be divided into six categories, including 5’UTR, 3’UTR, exon, intergenic, intron, and promoter. The peaks of H3K14la covered a large proportion of exons (40.64%) and promoters (37.91%), while only 12.37% were found in the intron regions (Figure S11A). Similar to the distribution of the peaks of H3K14la, H4K12la was also found mainly in exons (50.68%) and promoters (34.89%), respectively (Figure S11B). We performed an overlap analysis on the genes regulated by H3K14la and H4K12la from the three biological replicates, 195 genes were associated with H3K14la and 329 genes with H4K12la (Figure S11C and D). In addition, KEGG enrichment analysis showed that genes regulated by H3K14la were enriched in the necroptosis pathway (Figure S11E and Table S18). Genes associated with H4K12la were mainly enriched in cell cycle, ribosome biogenesis, phosphatidylinositol signaling system, and autophagy pathway (Figure S11F and Table S19). The overlapping analysis also revealed that 281 and 147 specific genes were modulated by H3K14la and H4K12la, respectively (Figure S12A). TSS enrichment with the sequences associated with H3K14la and H4K12la showed that the sequences were more enriched around TSS (3 kb upstream of TSS) and TES (Transcription End Site) regions (Figure S12B and C). These results suggest that histone H4K12la and H3K14la may actively regulate gene transcription in *T. gondii*.

Notably, many specific genes associated with H3K14la code for proteins in kinesin complex, which are related to microtubule motor activity participating in microtubule movement pathway (Figure S13A and Table S20). And the proteins encoded by the specific genes associated with H4K12la were predominantly involved in ATP and protein binding, which are closely associated with neurotransmitter transport pathways (Figure S13B and Table S21). Furthermore, the genes regulated by both H3K14la and H4K12la were closely associated with the activity of a variety of enzymes, including ubiquitin-protein transferase, protein serine/threonine kinase, phosphotransferase, acid-amino acid ligase, participating in the cellular protein modification process and microtubule-based motor pathways (Figure S13C and Table S22).

### TgPFKII is an important regulator of lactylation

Recent study has discovered that phosphofructokinase II (PFKII) was often induced in a hypoxic microenvironment, which greatly contributes to enhanced glucose metabolism, leading to increased levels of pyruvate [38]. To localize the expression of native PFKII in *T. gondii*, His-tagged PFKII fusion protein was generated and immunized into animals to produce TgPFKII-specific antibody (Figure 6A, Figure S14A and B). The native TgPFKII, which has a molecular weight of approximately 131 kDa, was identified by Western blotting, and the expression of PFKII in *T. gondii* ME49 strain was significantly higher than that in *T. gondii* RH strain (Figure 6B and C). IFA results showed that TgPFKII was widely distributed in multiple discrete compartments in intracellular and extracellular stages (Figure S14C and D). In addition, a total of seven proteins were identified to interact with TgPFKII by immunoprecipitation test and mass spectrometry analysis (Table S23), including proteasome 26S regulatory subunit (*TGME49_313410*), uncharacterized protein (*TGME49_205320*),uncharacterized protein (*TGME49_240220*) alanine-tRNA ligase (*TGME49_219540*), hydrolase 4 domain-containing protein (*TGME49_286580*), elongation factor 2 family protein (*TGME49_286080*), and histidyl-tRNA synthetase (*TGME49_280600*). As shown in Figure S15A-D, these proteins were differentially modified by 2-hydroxyisobutyrylation, crotonylation, and lactylation in the two *T. gondii* strains (Table S24), also in consistence with our previous findings [15].

**Figure 6.**
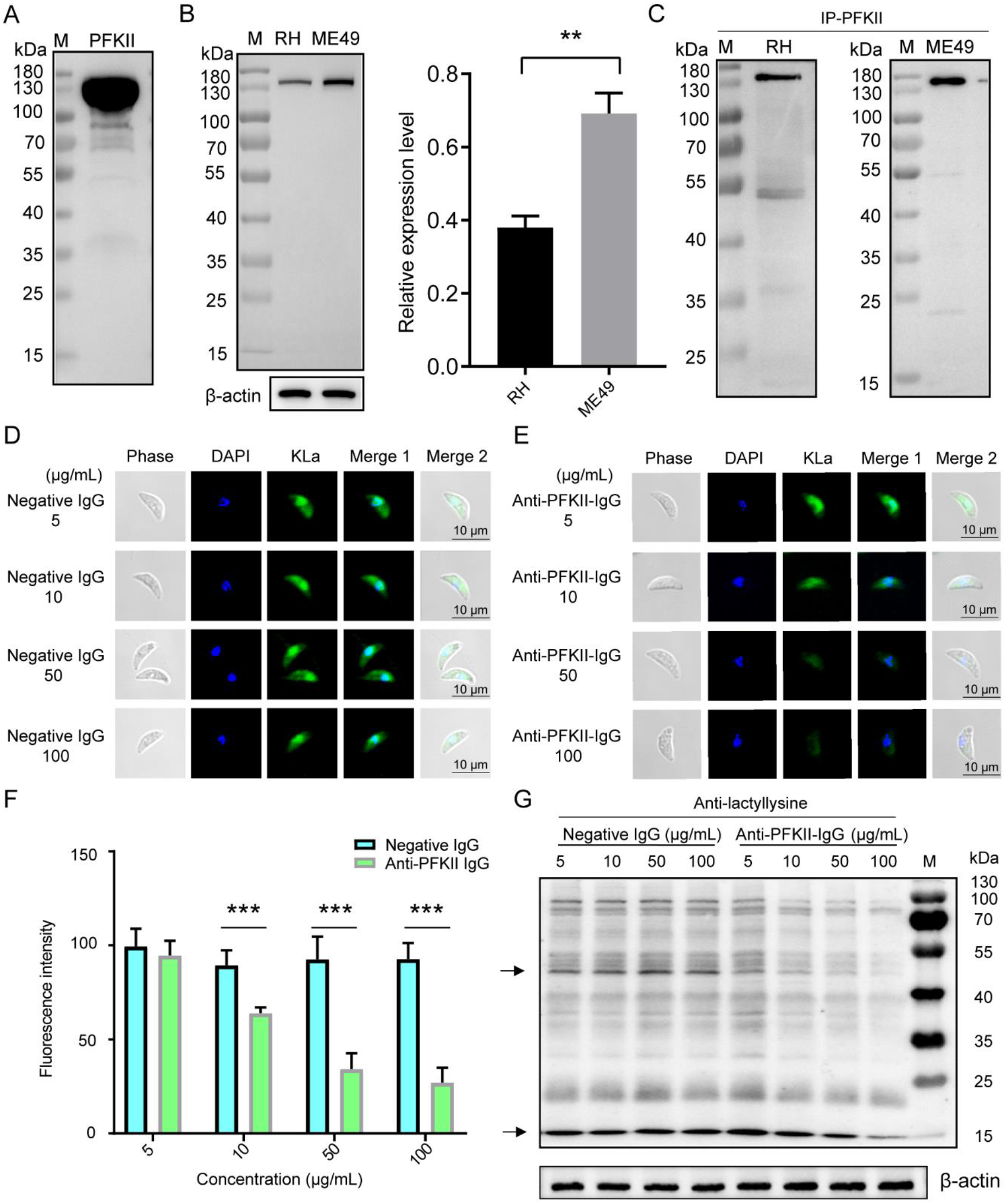
Anti-TgPFKII antibodies reduced protein lactylation. **A.** The purified recombinant protein TgPFKII was verified by Western blotting. **B.** Western blotting analysis of the expression levels of TgPFKII (131 kDa) in *T. gondii* RH and ME49 strains. β-actin was used for normalization. The data are presented as the mean ± SEM of three independent experiments (\**P* < 0.05, \*\**P* < 0.01, Student’s t-test). **C.** The soluble proteins obtained from both the *T. gondii* RH and ME49 strains was immunoprecipitated with anti-PFKII antibody, and the products were detected by Western blotting with anti-PFKII antibody. **D.** Negative group means that different concentrations of negative IgG (Mouse IgG) were co-cultured with *T. gondii* and analyzed for lactylation level (green) by IFA analysis. **E.** Different concentrations of anti-TgPFKII antibodies were co-cultured with *T. gondii* and the level of lactylation was analyzed by IFA analysis (green). **F**. Immunofluorescence intensity statistics were performed for the experimental (anti-PFKII antibodies) and negative groups, and significance analysis was performed using Student’s t test. All data are shown as mean and standard deviation (N **>** 10). \*\*\**P* < 0.001 versus control. **G.** Western blotting assay was used to test the effect of different concentrations of anti-PFKII antibodies on the lactylation level of *T. gondii* RH strain. β-actin was used for normalization. PFKII, phosphofructokinase.

The effect of TgPFKII on protein lactylation was confirmed by addition of TgPFKII-specific antibodies to the parasite culture as previously described [39]. The antibodies were able to infiltrate free tachyzoites and broadly disseminated in the cytoplasm illustrated by IFA and 3D Laser Scan Technology (Figure S16A, Movie 1 and 2). And the anti-TgPFKII antibody inhibited lactylation in a dose dependent manner (Figure 6D-G), suggesting that TgPFKII is an important component in the pathway of protein lactylation *in T. gondii* (Figure 6G). However, the anti-PFKII antibody had no effect on both *Plasmodium* (trophozoite) and *Trypanosoma*, though it strongly reduced lactylation level in Vero cells (Figure S16B-D).

### Antibodies to histone deacetylases and histone lysine acetyltransferase imposed significant effect on their enzymatic activity

Histone deacetylases (HDACs) and histone lysine acetyltransferase (MYST-A) control the acetylation status of histones in association with gene expression and downstream biological events [40] [41]. In *T. gondii*, five genes coding for HDAC class I/II including TgHDAC1, TgHDAC2, TgHDAC3, TgHDAC4, and TgHDAC5 have been identified, and which are potentially involved in histone PTMs [42]. The enzymes contain various functional domains. TgHDAC3 contains a PTZ00063 superfamily domain; TgHDAC2 contains a class I histone deacetylase structural domain (PRK12323 superfamily); TgHDAC4 contains arginase-like and histone-like hydrolases and trypanosomal VSG structural domain, and TgMYST-A contains a PLN00104 superfamily domain (Figure 7A). To localize the expression of the four proteins (TgHDAC2, TgHDAC3, TgHDAC4, and TgMYST-A) in *T. gondii*, we generated specific antibodies, which were verified by Western blotting (Figure 7B, Figure S17A and B). TgHDAC3 and TgMYST-A were located in the cytosol and nuclei in the parasites of both extracellular and intracellular phases (Figure 7C and Figure S17C), as reported in previous studies [41, 43, 44]. Notably, TgHDAC3 was distributed more in the cytosol than the nuclei in tachyzoites. In addition, TgHDAC2 and TgHDAC4 were also located in the cytoplasm and nucleus.

**Figure 7.**
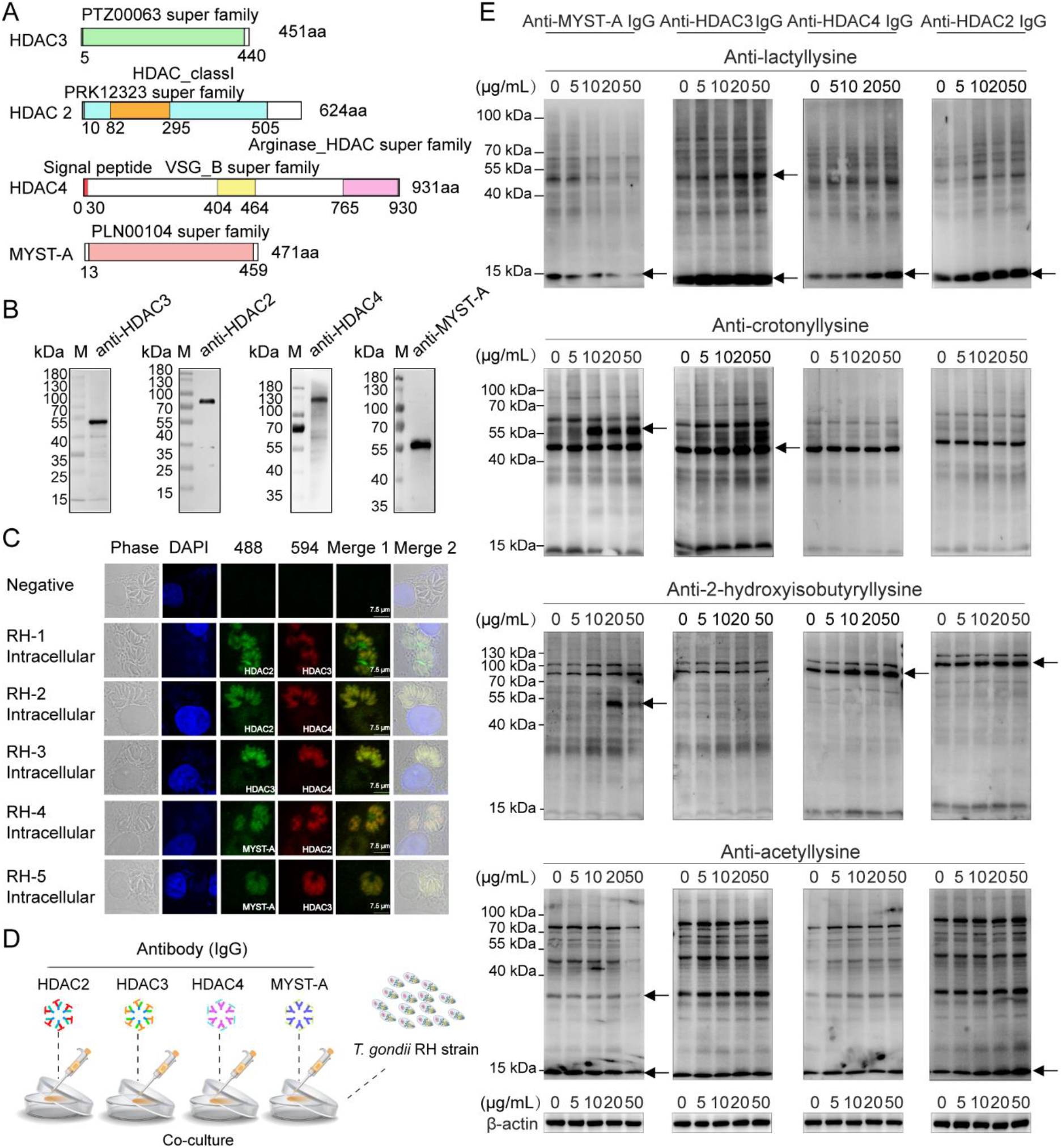
Preliminary analysis of the function of TgHDAC2, TgHDAC3, TgHDAC4, and TgMYST-A in associated with protein lactylation. **A**. Sequence characteristics of TgHDAC2, TgHDAC3, TgHDAC4, and TgMYST-A. **B**. Western blotting analysis of native TgHDAC2 (71 kDa), TgHDAC3 (53 kDa), TgHDAC4 (101 kDa), and TgMYST-A (56 kDa) expression level by TgHDAC2, TgHDAC3, TgHDAC4, and TgMYST-A specific antibodies. **C.** Indirect immunofluorescence assays (co-localization) of TgHDAC2, TgHDAC3, TgHDAC4, and TgMYST-A. **D.** Flow chart for the effect of specific antibodies on PTMs. **E.** Modulation of anti-TgHDAC2, anti-TgHDAC3, anti-TgHDAC4, and anti-TgMYST-A antibodies on lactylation, 2-hydroxyisobutyrylation, crotonylation, and acetylation of *T. gondii*. The difference of protein modification level was indicated by arrows. β-actin was used for normalization. HDAC2, HADC3, HDAC4, histone deacetylase; MYST-A, histone lysine acetyltransferase.

We then explored whether the four enzymes had an effect on PTMs in tachyzoites in vitro culture (Figure 7D). Anti-TgHDAC2, anti-TgHDAC3, and TgHDAC4 antibodies increased the overall level of protein lactylation, suggesting that these enzymes negatively regulated protein lactylation in *T. gondii*. And anti-TgHDAC3 antibody enhanced protein crotonylation, while anti-TgHDAC4 and anti-TgHDAC2 antibodies have no such effect. In addition, anti-TgHDAC2 and anti-TgHDAC4 antibodies increased 2-hydroxyisobutyrylation of non-histone proteins. Furthermore, we also observed that addition of anti-TgHDAC2 antibody increased overall protein acetylation, and anti-TgHDAC3 only slightly increased non-histone protein acetylation. By contrast, anti-TgMYST-A antibody could reduce protein lactylation and acetylation. Notably, when the concentration of the anti-TgMYST-A antibody was greater than 20 μg/mL, the levels of protein 2-hydroxyisobutylation and crotonylation of non-histone proteins (about 55 kDa) significantly increased (Figure 7E).

We also examined whether TgHDAC2, TgHDAC3, TgHDAC4, and TgMYST-A affected the modification of the H3K14la and H4K12la as well as the expression of TATA proteins by Western blotting. The results revealed that anti-TgHDAC3 and anti-TgHDAC4 antibodies slightly increased histone H3K14la level (Figure S17D). On the other hand, lactylation on H4K12 was increased by anti-TgHDAC2 and anti-TgHDAC3 antibodies, while the level of H4K12la remained unchanged by anti-TgHDAC4 and anti-TgMYST-A antibodies. And anti-TgMYST-A antibody significantly enhanced the expression level of TATA-binding proteins (Figure S17D).

## Discussion

As an intracellular parasite, *T. gondii* utilizes multiple carbon sources to ensure its own energy demands, approximately 80% of internalized glucose in *T. gondii* is catabolized to lactate at the tachyzoite stage [28, 29, 45], and the intermediates and end products of its metabolic processes are recognized to act as essential signals, which modulate cellular activity and gene expression [46, 47]. Lactate was initially regarded as a metabolite of glycolysis and has recently been considered as a signal that stimulates protein lactylation [20]. In this study, an atlas of the lactylome of the *T. gondii* RH strain was established for the first time, and this atlas contained 955 proteins with 1964 lactylation sites. The flux of lactylation was close to crotonylation, which was the second only to phosphorylation and 2-hydroxyisobutyrylation (Table S25). Our studies revealed that lactylated proteins were distributed in diverse compartments (especially the mitochondria) and were significantly enriched in energy metabolism processes such as oxidative phosphorylation, glyoxylate and dicarboxylate metabolism, glycolysis/gluconeogenesis, and TCA cycle (Figure 2, Figure S2 and S3). Our results are also in line with previous findings in *Trypanosoma brucei* and rice, where lactylation is important for gene transcription and central carbon metabolism [48, 49]. We therefore focused on the link between lactylation and the processes of energy metabolism. We found that protein lactylation in the high glucose culture environment was significantly higher than that in the low glucose medium (Figure S18A). Further, inhibition of lactate generation with sodium dichloroacetate and oxamate resulted in less histone Kla and inhibition of mitochondrial respiratory chain complex I with rotenone increased the level of histone Kla (Figure S18B–D), indicating that protein lactylation is positively correlated with glycolysis and mitochondrial metabolism in *T. gondii*. Moreover, we identified diverse lactylated rate-limiting enzymes involved in energy metabolism processes, including HK, PYK, and CSI. Interestingly, some glycolytic enzymes in *T. gondii* contain two isoforms that exhibit differential modification patterns at both the tachyzoite and bradyzoite stages [3, 50]. Three lactylation sites (K109, K136, and K4) on PYK1 and two lactylation sites (K328 and K336) on PYK2 were identified.

In addition, lactate dehydrogenase is an important enzyme for the conversion of pyruvate to lactate, and *T. gondii* expresses LDH1 only at the tachyzoite stage [51]. Here, we only identified LDH1 with five lactylation sites (K43, K94, K211, K218, and K322) in the *T. gondii* RH strain, and these sites might be essential for balancing the levels of pyruvate and lactate. However, the biological significance of the differential modifications remains to be further investigated. Under eutrophic conditions, oxidative phosphorylation in mitochondria is the main pathway for ATP synthesis and is sufficient to support *T. gondii* reproduction. However, under physiological conditions, when ATP production is not sufficient, LDHs-mediated lactate fermentation is essential for the energy supply [3]. Thus, lactylation may have a more important role in regulation of energy metabolism in *T. gondii* under physiological condition. Earlier studies reported that phosphorylation was mainly associated with signaling transduction in *Toxoplasma* [52]; palmitoylation was more importantly linked to the process of *Toxoplasma* invasion of host cells [9]; ubiquitination was mainly associated with the regulation of the cell cycle transition of *T. gondii* [19]. In contrast to these PTMs, the lactylation of *T. gondii* is highly correlated with energy metabolic processes.

In animal cells, glycolytic/glycolytic fluxes are highly regulated by fructose 2,6-diphosphate levels, and fructose-2-phosphate kinase mediates the formation of fructose-2,6-diphosphate converted from fructose-6-phosphate [53, 54]. The level of lactylation dropped dramatically after anti-TgPFKII antibody treatment in *T. gondii* and Vero cells, whereas it has no effect on both *Plasmodium* and *Trypanosoma* species (Figure 6 and S16), implying the effect of enzyme-specific antibodies is parasite dependent. In our previous study, TgPFKII was the regulatory enzyme containing the highest number of sites in glycolysis pathway of *T. gondii* [15]. Here, we further confirmed that TgPFKII is crotonylated and 2-hydroxyisobutyrylated (Figure S19), implying that 2-hydroxyisobutyrylation and crotonylation may modulate the activity of TgPFKII and protein lactylation.

There is growing evidence that enzymes, like HDACs and MYST-A, are involved in the regulation of various cell functions [44, 55]. In this study, we revealed that TgHDAC3 can act as delactylase, decrotonylase, and deacetylase in tachyzoites. And TgHDAC2 is capable of regulating lactylation, acetylation, and histone 2-hydroxybutyrylation. We further demonstrated that TgHDAC4 had both delactylase and de-2-hydroxymethylase activities in tachyzoites. Notably, TgHDAC4 contains not only the Arginase HDAC superfamily structural domain, but also the VSG structural domain. The main function of the VSG structural domain is to assist the parasite in immune evasion [56, 57]. Unlike the other HDACs, only TgHDAC4 contains a signal peptide (0-26 aa) and therefore it may be a secreted protein that not only functions as a modifying regulatory enzyme, but also might be a molecule associated with interaction with host immune systems. Previous studies have shown that HDACs has effective delactylase activity [44, 58], which was also confirmed in this study. In this study, TgMYST-A, as a histone lysine acetyl transferase, was found to be able to improve histone acetylation and lactylation level, which might have a synergetic effect on the differentiation of *T. gondii*. Notably, anti-TgMYST-A antibody could increase the levels of crotonylation and 2-hydroxyisobutyrylation on non-histone proteins (55 kDa), a subject worth of further investigation.

Histone is the primary structural component of chromatin, and histone PTMs play an indispensable role in the epigenetic regulation of eukaryotic genes [35]. As shown in Figure 4, 17 Kla sites were detected on three core histones (H2Bb, H3, and H4) and two variant histones (H2A.Z and H2B.Z), which were mainly located in the N-terminal tail domains. Interestingly, most histone sites could undergo multiple PTMs rather than a single PTM. Notably, the histones H2BbK46 and H2BbK98 were monomodified in the globular domain, which might represent specific regulation of gene transcription associated with energy metabolism in *T. gondii*. In human and mouse cells, 28 Kla sites have been identified on core histones, which are closely associated to gene activation [22]. We found that the histone Kla sites of *T. gondii* are in certain circumstance similar to that of human and mouse, which may also critically regulate gene transcription. Among all 193 PTMs sites, 69 (35.8%) and 85 (43.5%) were distributed in horizontal line and globular domain, respectively, suggesting that PTMs co-regulate genes by altering the interactions among histones and DNA. Study reported that H4k12la is enriched at the promoters of glycolytic genes and activates transcription in Aβ plaque-adjacent microglia, thereby increases glycolytic activity [59]. The ChIP-data indicated that the binding regions of H4K12la and H3K14la on the chromosomes were concentrated in the vicinity of TSS and were associated with cellular protein modification and microtubule motor process in *T. gondii*. This is the first report to reveal the important relationship and function between H4K12la and H3K14la in *T. gondii* tachyzoites.

PTM crosstalk is important for regulation of biological processes [60–62]. Here, we found that multiple PTMs occur on the same lysine of the same protein, for example 392 lysine sites were commonly modified by lactylation (Kla), crotonylation (Kcr), and 2-hydroxyisobutyrylation (Khib) (Figure S20, Table S26 and S27). And the Kcr, Khib, and Kla proteins were mainly distributed in mitochondria, associated with ligase activity, and significantly enriched in processes such as ribosomal RNA monophosphate biosynthesis, tRNA metabolism, and protein translation (Figure S21 and Table S28). Furthermore, three PTMs were related to myosin head-motor domain, implying that these PTMs may functionally regulate the proteins in host cell invasion by *T. gondii* (Figure S22A and Table S29). Additionally, Kcr, Khib, and Kla proteins were enriched in glycolysis/gluconeogenesis, propanoate metabolism, HIF-1 signaling, the citrate cycle, aminoacyl-tRNA biosynthesis, RNA transport, oxidative phosphorylation, and ribosome processing (Figure S22B and Table S30). Therefore, multiple PTMs may jointly regulate various biological processes.

In conclusion, we established the first comprehensive atlas of protein lactylation in the zoonotic parasite *T. gondii* and revealed that lactylation profoundly modulate glucose and genes activation in various biological processes in the parasite (Figure S23). The data provided new front in digging anti-parasite drugs.

## Materials and methods

### Experimental design and statistical rationale

The global Kla of total proteins derived from the *T. gondii* RH strain was analyzed by label-free method. Three biological replicates of protein lactylation were analyzed to evaluate the accuracy of the data obtained in the experiment (Table S1). A qualitative analysis of lactylation was performed, and Fisher’s exact test was used to determine whether the data were statistically significant. R scripts were used for the statistical tests and pattern generation.

### Experimental animals

Female Kunming mice (6–8 weeks old) and rats (200 g) were purchased from Liaoning Changsheng Biological Technology Company in China.

### Cultivation of *T. gondii*

*Toxoplasma gondii* tachyzoites were cultured in a confluent monolayer of monkey kidney adherent epithelial (Vero) cells and incubated at 37°C with 5% CO_2_ in high sugar medium and low sugar medium (Catalog No. SC102-01 and SC104-01, Seven Biotech, Beijing, China) containing fetal bovine serum (Catalog No. 10099, Thermo Scientific, Waltham, MA) as previously described [63, 64]

### Parasite purification and protein extraction

Tachyzoites from the peritoneal fluid of *T. gondii* RH strain infected mice were filtered through 5.0-μm nucleopore filters and then collected by centrifugation into a centrifuge tube (No. CFT920150, Jet Bio-Filtration, Guangzhou, China). The Percoll (Catalog No. 17-0891-09, GE Healthcare, Sweden) gradient centrifugation method was used for separation and purification. The purified parasite cells were washed with cold phosphate-buffered saline (PBS-1×) and sonicated in the presence of a cocktail of protease inhibitors (Catalog No. HY-K0010, MCE, New Jersey) three times in lysis buffer (1% protease inhibitor mixture and 8 M urea) using a high-intensity sonicator (Catalog No. 01A566, Scientz, Ningbo, China). The lysate was centrifuged at 12,000 g (4 °C) for 12 minutes to remove insoluble cell fragments, and the sample concentration was measured using a BCA kit (Catalog No. P0010, Beyotime, Shanghai, China).

### Trypsin digestion of parasite proteins and high-performance liquid chromatography fractionation

The protein solutions were reduced at 56 °C for 30 minutes using 5 mM dithiothreitol and alkylated at 37 °C for 15 minutes using 11 mM iodoacetamide in a dark. Subsequently, the protein samples were diluted with 100 mM tetraethylammonium bromide, and trypsin was added for digestion overnight. The peptides were separated by high pH reversed-phase separation (RPF) using a Thermo Betasil C18 column.

### Antibody-based PTM enrichment and LC-MS/MS analysis

The peptides were dissolved in an immunoprecipitation (IP) buffer solution (100 mM NaCl, 1mM EDTA, 50 mM Tris-HCl, 0.5% NP-40, pH 8.0), and the supernatant was transferred to the pre-washed panlysine-lactylation antibody conjugated beads (Catalog No. PTM-1401, PTM BIO, Hangzhou, China) and incubated overnight at 4 °C. The peptides bonds on the beads were eluted with 0.1% trifluoroacetic acid and vacuum dried. The resulting peptides were desalted using C18 ZipTip (Catalog No. ZTC18S096, Millipore, Boston, MA) as indicated previously described [22]. The rest of the steps were set as described previously [15].

### Database search

The secondary mass spectrometry data were retrieved using MaxQuant (v1.5.2.8) software from the *ToxoDB* 46.0 database (8,322 sequences, https://toxodb.org/toxo/), and the false positive rate (FDR) caused by random matching through the anti-database was calculated. The rest of the steps were set as described previously [15].

### Preparation of specific antibodies

The genes coding for *T. gondii* histone deacetylase HDAC2 (gene ID, *TGME49_249620*),HADC3 (gene ID, *TGME49_257790*), HDAC4 (gene ID, *TGME49_227290),* histone lysine acetyltransferase MYST-A (gene ID, *TGME49_318330*) and phosphofructokinase PFKII (gene ID, *TGME49_ 226960*) were amplified from the genome of *T. gondii* RH strain by polymerase chain reaction (PCR) with gene-specific primers (Table S31). The PCR products were cloned into the pET-28a vector and pGEX-4T-1 vector, respectively. The constructs containing the target genes were expressed in *Escherichia coli* BL21 (DE3) cells (Catalog No. CD601-02, TransGen Biotech, Beijing, China), and the His-tagged and GST-tagged recombinant proteins were purified according to the manufacturer’s instructions (Catalog No. CW0010S, CWBIO, Beijing, China) and evaluated by SDS-PAGE and Western blotting. Rats and mice (n = 10) were subcutaneously immunized 4 times with His-tagged recombinant proteins emulsified with Freund’s adjuvants. Protein-specific antibodies were verified with GST-tagged recombinant proteins. IgG were purified from the immune sera (mouse/rat) using Protein A (Catalog No. 17061801, Cytiva, JVIA-W) and Protein G Sepharose 4 Fast Flow Resin (Catalog No.17075601, Cytiva, JVIA-W), respectively.

### Western blotting

*Toxoplasma gondii* lysate (20 μg) was dissolved in polyacrylamide gel electrophoresis, and the proteins were transferred to a polyvinylidene fluoride membrane (Catalog No. 162-0177, Bio-Rad, CAL). The membrane was blocked with 5% skimmed milk at 37°C for 45 minutes and then incubated at overnight 4°C with an anti-lactyllysine antibody (Catalog No. PTM-1401, PTM BIO, Hangzhou, China). The membrane was washed four times with phosphate-buffered saline (PBS-1×) buffer and incubated with a secondary antibody at 37°C for 50 minutes (Catalog No. 31430, Thermo Scientific™ Pierce, Waltham, MA). The procedure for the Western blotting assays of the recombinant proteins are the same as above (antibody dilution ratio of 1: 200). Anti-TATA binding protein and anti-beta tubulin antibodies were purchased from PTM bio (Catalog No. PTM-5007 and PTM-5028, PTM BIO, Hangzhou, China).

### Immunofluorescence assay

The tachyzoites were fixed on glass slides with prechilled paraformaldehyde for 10 minutes. After permeabilization with 0.1% Triton X-100 for 15 minutes, the parasites were then blocked with 5% skimmed milk for 1 h and then incubated with an anti-lactyllysine primary antibody (Catalog No. PTM-1401, PTM BIO, Hangzhou, China) that recognizes lactyllysine for 1 h. The slides were subsequently incubated with Alexa Fluor secondary antibody (Catalog No. 31430 and 31460, Thermo Scientific™ Pierce, Waltham, MA) at 37°C for 30 minutes, and the nuclei were stained as described previously [65, 66]. The negative control was the tachyzoites incubated with negative rabbit or mouse IgG (Catalog No. AG-0021 and AG-0051, Dingguo Biotechnology, Beijing, China). Parasites were stained with ProLong Gold antifade mountant (DAPI) (Catalog No. P36982, Invitrogen, CA). High-resolution images were then captured by a confocal microscopy (Catalog No. TCS-SP8, Leica, Wetzlar, Germany). IFA methods for TgHDAC2, TgHDAC3, TgHDAC4, TgMYST-A and TgPFKII as above.

To verify the permeability of antibodies into live *T. gondii*, anti-TgPFKII antibodies (1:200) was added to the medium and incubated with *T. gondii* for 3–4 h and then fixed in cold methanol. The tachyzoites were permeabilized by 0.1% Triton X-100 for 20 minutes and then washed twice with PBS. The Alexa Fluor 488 secondary antibodies were added directly at 37°C for 40 minutes. The glass slide was washed three times with PBS-1 × buffer and the nuclei were stained as described previously [65].

### Inhibition assay with anti-TgHDAC2, anti-TgHDAC3, anti-TgHDAC4, anti-TgMYST-A, and anti-TgPFKII antibodies

The specific antibodies of TgHDAC2, TgHDAC3, TgHDAC4, TgMYST-A, and TgPFKII were tested for their effects on the levels of crotonylation, 2-hydroxyisobutyrylation, lactylation, and acetylation. Before complete release of *T. gondii* RH tachyzoites from Vero cells, antibodies of TgHDAC2, TgHDAC3, TgHDAC4, TgMYST-A or TgPFKII were added at concentrations of 0, 5, 10, 20, and 50 μg/mL, respectively, to the cells at 37 °C for 4 h. The tachyzoites of *T. gondii* RH were centrifuged and rinsed with cold sterile PBS-1 **×** buffer for 2 h. Total protein was extracted using a protein extraction kit (Catalog No. FNN0011, Thermo Fisher Scientific, Waltham, MA). The effect of different concentrations of antibodies on PTMs was examined by Western blotting. Mouse β-actin (Catalog No. PTM-5018, PTM BIO, Hangzhou, China) was used for normalization.

### Immunoprecipitation experiment for TgPFKII

In order to identify the proteins interacting with TgPFKII, the tachyzoites of *T. gondii* were collected and purified, sonicated 3 times for 5 s in the lysate buffer (Catalog No. C500005, Sangon Biotech, Shanghai, China) containing protease inhibitors. Protein concentrations were determined by the BCA kit to ensure that the protein masses of the *T. gondii* RH and ME49 strains were equal, while blank and negative controls were also set up. Anti-TgPFKII antibody (2 μg) was added to the cell lysate and an equivalent amount of irrelevant IgG was used as a negative control. The samples were mixed with overnight agitation at 4°C. Then 20 μL Protein A or Protein G beads were added and mixed for 1-3 h at 4°C. The beads were precipitated by centrifugation at 4°C for 30 seconds, followed by washing 5 times with 500 μL 1 × lysis buffer. Finally, the sediment was heated at 100°C for 10 minutes after being mixed with 20-40μL 5× SDS sample buffer. The beads were precipitated by centrifugation and the supernatants were subjected to Western blotting experiments with lactyllysine, 2-hydroxybutyryllysine and crotonyllysine antibodies (Catalog No. PTM-801 and PTM-502, PTM BIO, Hangzhou, China). The immunoprecipitated proteins were dissolved in SDS-PAGE gel. Protein bands were excised and analyzed by mass spectrometry using LC-MS/MS on Thermo Q exclusive HF as previously described [67].

### Lactate metabolic inhibitors on histone lactylation

Before complete release of *T. gondii* RH tachyzoites from Vero cells, inhibitors were added at concentrations of 0, 5, and 20 mM, respectively, to the cells at 37 °C for 4 h. After 4 h, tachyzoites of *T. gondii* RH were centrifuged and rinsed with cold sterile PBS-1× buffer after 2 h. Total protein was extracted and the lactylation level of *T. gondii* RH strain was examined by Western blotting analysis. Sodium dichloroacetate (DCA), sodium oxalate and rotenone were purchased from Selleck (Catalog No. S8615 S6871, and S2348, Selleck, Houston, TX).

### Chromatin Immunoprecipitation Sequencing

ChIP experiments were performed using histone anti-H4K12la and anti-H3K14la antibodies (Catalog No. PTM-1411 and PTM-1414, PTM BIO, Hangzhou, China) respectively. *T. gondii* parasites were collected in centrifuge tube and washed with 1×PBS buffer. After centrifugation and removing the supernatant, 1% formaldehyde in PBS buffer was added to cross-link DNA and proteins for about 10 minutes at 37°C. Glycine was added to stop the cross-linking reaction. PBS containing 0.5% bovine serum and protease inhibitor was then added to wash the parasites. The samples were collected by centrifugation and resuspended in 200 μL lysis buffer (1% SDS, 10 mM EDTA, 50 mM Tris-HCl, pH 7.5) and incubated on ice for 10 minutes. The lysate was sonicated and one tenth was removed as an input control. The remaining lysate was immune-precipitated by adding 4 μg of either anti-H4K12la or anti-H3K14la antibodies to ChIP dilution buffer (1% Triton, 150 mM NaCl, 2 mM EDTA, 20 mM Tris-HCl, pH 7.5), which had been pre-incubated with protein A/G magnetic beads (Catalog No. 10003D, Invitrogen, Carlsbad, CA). The mixture was incubated overnight at 4°C and then washed twice with RIPA buffer (1 mM EDTA, 10 mM Tris, 0.1% SDS, 0.1% Triton X-100, 0. 1% Na-deoxycholate), LiCl buffer (0.5% NP-40, 0.25 M LiCl, 0.5% Na-deoxycholate), and Tris-EDTA buffer, respectively. The DNA fragments were eluted with elution buffer (1% SDS, 0.1 M NaHCO3) and purified by phenol-chloroform extraction and precipitated with glycogen and ethanol. The ChIP-DNA fragments were used to construct libraries using the Illumina protocol, which were then sequenced by the Illumina sequencing platform (Illumina NovaSeq 6000, Illumina, San Diego, CA).

### Raw data filtration of ChIP

Trimming of illumina paired-end and single-end data (low quality bases) was performed by the Trimmomatic software (version 0.36). It works in conjunction with FASTQ or gzipp’ed FASTQ. With “simple” trimming, each adapter sequence is tested against the read, and if a sufficiently accurate match is detected, the read is trimmed appropriately.

### Sequence alignment

Clean reads were mapped to the genome of *T. gondii* using Bowtie2 (version. 2.3.4.3) with default parameters. The alignment file (SAM) was transformed to BAM file and was filtered by the following criteria: (1) only keep unique aligned reads; (2) remove reads with low mapping qualities (< 30).

### Peak calling

The MACS2 (version: 2.1.2) was used to identify transcription factor binding sites and analysis genomic complexity to assess the significance of ChIP-enriched regions.

### Peak annotation

The peaks were annotated by ChIPseeker (version: 2.16.0), and an R package for annotating ChIP-seq data analysis. The software supports both comparative analysis of ChIP peak maps and annotations. Peaks from three experiments that were annotated with the same gene were combined into the final identified peak.

### GO annotation

The lactylated proteins were subjected for Gene Ontology (GO). The process of GO annotation mainly converts protein IDs into UniProt IDs matched with GO IDs and then retrieves the corresponding information from the UniProt-GOA database.

### Domain annotation

Using InterProScan software and the corresponding InterPro domain database, the identified proteins were annotated with protein domains.

### KEGG pathway annotation

The proteins were annotated using KAAS tool and matched to the corresponding pathways using the KEGG mapper.

### KOG annotation

The sequence of modified proteins was matched using the version 2.2.28 of the KOG database, and protein annotation information was obtained using the basic local alignment search tool.

### Subcellular localization

The WoLF PSORT software performs subcellular location annotation of the submitted proteins.

### Motif analysis

The compositional characteristics of the 10 amino acids upstream and downstream of the lactylation site were analyzed by MoMo Software. The conditions for qualifying for motif are a number of peptides containing a particular sequence form greater than 20 and *P* < 0.000001. The degree of variation (DS) in the frequency of occurrence of amino acids near the modification site is shown by the heat map. DS, difference score.

### GO, KEGG, and protein domain enrichment analyses

For every group, a two-tailed Fisher’s exact test (*P* value < 0.05) was used to assess the statistical significance of the results from the enrichment analyses.

### Protein interaction network analysis

The gene numbers of the all modified proteins were matched against the STRING (version 11.0) database to obtain protein interaction relationships (high confidence >0.7). The protein interaction network was visualized using the R package “networkD3”.

---

Movie S1. The distribution of anti-TgPFKII antibody was detected by 3D Laser Scan Technology (Sample 1)

Movie S2. The distribution of anti-TgPFKII antibody was detected by 3D Laser Scan Technology (Sample 2, Negative)

## Ethics statement

All animal manipulations performed in this study were carried out in accordance with the Shenyang Agricultural University Animal Husbandry Guidelines. The Ethics Committee of Shenyang Agricultural University approved the experimental animal experiments (Permit No. SYXK<Liao>2020-0001).

## CRediT authorship contribution statement

**Qijun Chen**: Conceptualization, Methodology, Data curation, Writing-Review & Editing, Supervision. **Deqi Yin**: Writing-Original draft preparation, Investigation, Validation, Data curation. **Ning Jiang**: Validation, Supervision. **Chang Cheng**: Validation. **Xiaoyu Sang**: Software. **Ying Feng**: Formal analysis. **Ran Chen**: Software.

## Data availability

The MS proteomics data have been deposited to the ProteomeXchange Consortium (http://proteomecentral.proteomexchange.org) via the PRIDE partner repository with the data set identifier PXD022700. The ChIP-seq data have been made available at the Sequence Read Archive (SRA) database under the BioProject accession number PRJNA791485 (https://submit.ncbi.nlm.nih.gov/subs/bioproject/).

ChIP-seq data (GSA: CRA006903) have been deposited in the Genome Sequence Archive at the National Genomics Data Center, Chinese National Center for Bioinformation, and are available via the https://ngdc.cncb.ac.cn/gsa. The data (proteomics data) reported in this paper have been deposited in the OMIX, China National Center for Bioinformation/Beijing Institute of Genomics, Chinese Academy of Sciences (https://ngdc.cncb.ac.cn/omix: accession no. OMIX001139).

## Acknowledgments

This study was supported by the grants from the National Key Research and Development Program of China (Grant Numbers 2017YFD0500400, 2017YFD0501200); the National Natural Science Foundation of China (Grant Numbers 31972702, 81772219); and CAMS Innovation Fund for Medical Sciences (CIFMS) [2019-I2M-5-042].

We appreciate very much for the technical and bioinformatic assistance from the scientists of PTM-bio, Hangzhou, China, and scientists at E-Gene for the technical assistance with ChIP-seq.

